# A stereotaxic atlas of primary cortical areas in the developing rat brain from postnatal days 8 to 20

**DOI:** 10.64898/2026.07.22.740117

**Authors:** Nicholas J. Sattler, Anna K. Grobengieser, James C. Dooley

**Author notes:** Corresponding author: James C. Dooley, Ph.D. Lead Contact: James C. Dooley. Author contributions: J.C.D. designed research; N.J.S. and J.C.D. performed research; N.J.S., A.K.G., and J.C.D. performed histology and microscopy; N.J.S., A.K.G., and J.C.D. analyzed data; N.J.S. and J.C.D. visualization; J.C.D writing – original draft; N.J.S., A.K.G., and J.C.D. writing – review & editing; J.C.D. supervision; J.C.D. resources.

## Abstract

Postnatal brain growth is non-linear, making precise stereotaxic targeting in the developing rat neocortex difficult without age-specific knowledge of cortical area locations. Traditional atlases visualize brain slices in the coronal plane, which can obscure top-down areal boundaries and sub-domains. To address these shortcomings, we created a developmental stereotaxic atlas that maps the neocortex of postnatal day (P) 8, P12, P16, and P20 in Sprague-Dawley and Long-Evans rats onto a coordinate grid. After using a stereotaxic device to create a grid of fluorescent probes, we extracted the brain, flattened neocortical tissue, and stained it for cytochrome oxidase, which is predominantly found in layer IV of primary cortical areas. Reconstructions of the primary somatosensory, auditory, and visual cortices demonstrate high structural reproducibility within each age and strain group. Our atlas captures the location of primary cortical areas at these 4 ages, showing that neocortical expansion is non-isometric, expanding preferentially along the rostral-caudal axis. Finally, we complement these top-down maps by extracting local neocortical surface angles from an existing coronal atlas, enabling proper electrode orientation to be tangential to the developing neocortex. Ultimately, this anatomically verified resource provides a standardized blueprint that eliminates resource-intensive trial-and-error mapping and maximizes experimental reproducibility in developmental systems neuroscience.

**Significance Statement:** Targeting specific neocortical areas in developing rats is uniquely challenging because non-linear brain expansion renders scaled adult coordinates inaccurate, while traditional coronal sections obscure top-down areal boundaries. To resolve this, we established a top-down stereotaxic atlas that maps primary sensory cortices onto flattened, cytochrome oxidase-stained tissue across early postnatal development (P8–P20) in both Sprague-Dawley and Long-Evans rats. By combining equidistant coordinate grids with empirical cortical surface angles, this resource provides an accurate, reproducible surgical blueprint. This reference tool eliminates trial-and-error coordinate mapping, reduces animal waste, and maximizes experimental precision for the developmental neuroscience community.

## Introduction

The rat has long been one of the most popular experimental organisms used in behavioral neuroscience, with the first comprehensive adult rat brain atlas published over 60 years ago (König and Klippel, 1963). These tools, along with stereotaxic devices, have enabled researchers to accurately target pharmacological injections and neurophysiological recordings in specific cortical areas and subcortical nuclei of the adult rat brain. More recently, methodological innovations have pushed the boundaries of developmental neuroscience, making neurophysiological recordings and stereotaxic injections in the neocortex of developing rats increasingly common (Dooley and van der Heijden, 2024). This technological shift has driven the need for atlases in developing rats like those found in adults. Because brain development is highly non-linear—particularly in the neocortex—simply ‘scaling down’ adult coordinates results in significant stereotaxic errors. Consequently, this has led individual labs to determine their own coordinates, resulting in unnecessary duplication that wastes both time and animal life.

To answer this need, atlases of the developing rat brain have been published for embryonic (Altman and Bayer, 1995) and postnatal ages (Zilles, 2012; Khazipov et al., 2015; Ramachandra and Subramanian, 2016; Chen et al., 2022; Bajic et al., 2023), some of which incorporate stereotaxic coordinates and histological staining. However, identifying and aiming for precise sub-regions within primary cortical areas is constrained by the traditional coronal perspective of existing atlases and a lack of distinctive structural markers in the developing neocortex. Without a top-down, overhead view of areal boundaries, navigating neocortical areas can remain challenging.

Here, we present a developmental neocortical atlas that maps stereotaxic coordinates directly onto flattened tissue stained for cytochrome oxidase (CO). CO readily labels layer IV of primary sensory areas from postnatal day (P) 5 through adulthood (Seelke et al., 2012). Reconstructions of stereotaxic coordinates on top of CO stained tissue are presented for rats at P8, P12, P16, and P20. These coordinates have been successfully used in several previous studies by our laboratory (Dooley and Blumberg, 2018; Reid et al., 2026) and are consistent with published values used by other labs that study the development of the rat neocortex.

## Methods

A total of 8 male and female Sprague-Dawley (6 M, 2 F) and 8 male and female Long-Evans (3 M, 5 F) Norway rats (*Rattus norvegicus*) were used from P8-P20, which included two rats per age. For each age and strain, the animal with the best histology quality was selected for the atlas. Pup sex, weights, and bregma-lambda distances can be seen in Table 1 and Figure 1. Pups were born to mothers housed in laboratory cages (48 x 20 x 26 cm) with a 12:12 light-dark schedule. Food and water were freely available. Expecting mothers were checked for pups daily, and the day of birth was considered P0. If litters consisted of more than eight pups, litters were culled down to eight pups (with equal numbers of males and females if possible) on or before P3 in order to ensure similar maternal care, reducing variability. The two animals at each age and strain always came from different litters. All experiments were conducted in accordance with the National Institutes of Health (NIH) Guide for the Care and Use of Laboratory Animals (NIH Publication No. 80–23) and were approved by the Institutional Animal Care and Use Committees of the University of Iowa and Purdue University.

**Table 1:** Bregma-lambda distance, weights, and sex for all included animals.

| *Age* | *B-L distance (mm)* | *Weight (g)* | *Sex* |
| --- | --- | --- | --- |
| Long-Evans |  |  |  |
| *P8* | 5.02 | 19.7 | F |
|  | 4.75 | 14.8 | M |
| *P12* | 5.51 | 27.7 | M |
|  | 5.46 | 25.4 | F |
| *P16* | 6.17 | 35.2 | F |
|  | 6.24 | 39.4 | F |
| *P20* | 7.32 | 52.1 | M |
|  | 7.08 | 53.5 | F |
| Sprague-Dawley |  |  |  |
| *P8* | 5.91 | 21.02 | M |
|  | 6.11 | 17.39 | F |
| *P12* | 6.78 | 31.45 | M |
|  | 6.52 | 30.5 | F |
| *P16* | 7.21 | 45.16 | M |
|  | 7.07 | 39.7 | M |
| *P20* | 7.52 | 61.3 | M |
|  | 7.41 | 58.3 | M |

**Figure 1.**
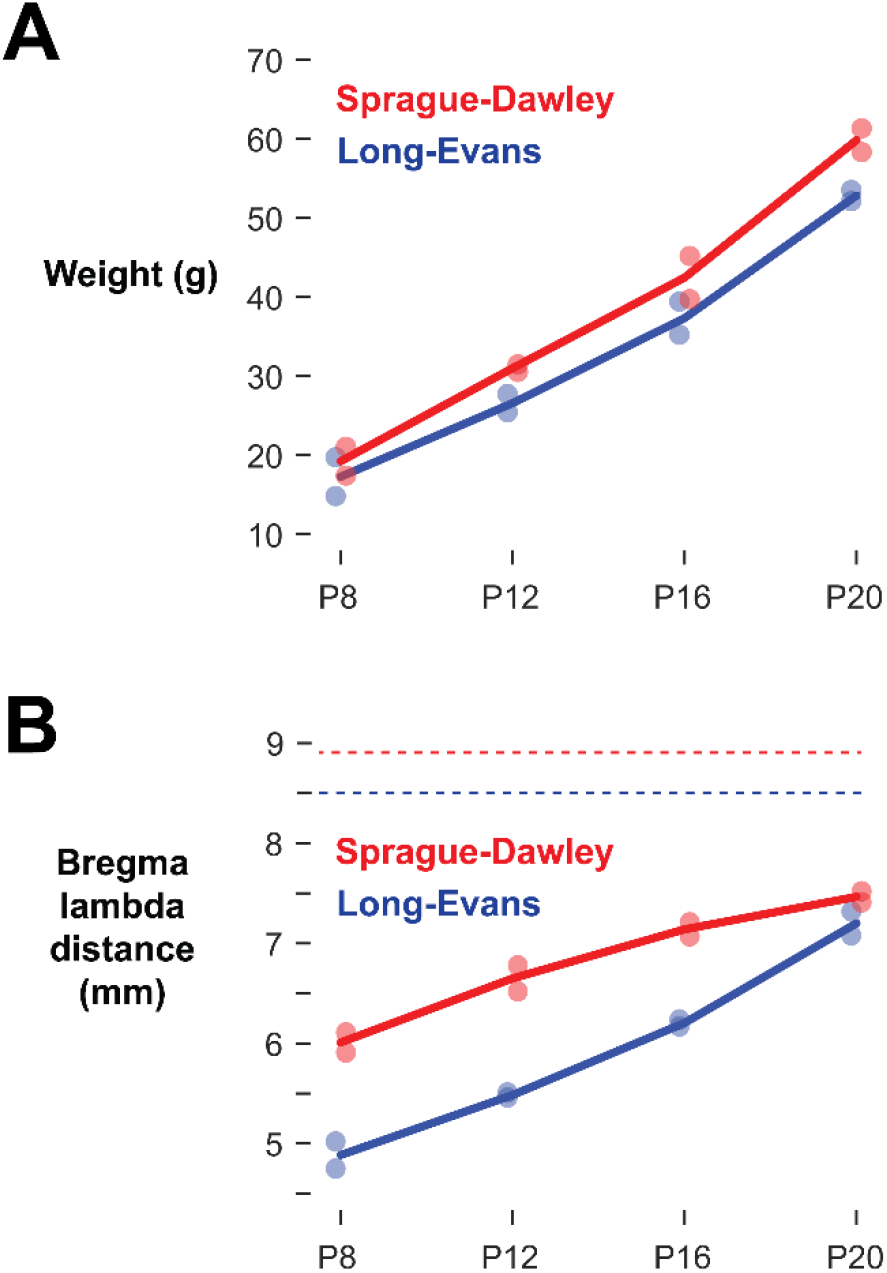
Postnatal body and skull growth across age by strain. **A** Total body weight (g) across early development. Colored dots indicate individual subject weights, and solid lines track the group mean. **B** Bregma-lambda distances (mm) across development. Solid lines connect early postnatal group means. Horizontal dotted lines at the top of the plot represent reference adult bregma-lambda distances for each strain (red: Sprague-Dawley; blue: Long-Evans), illustrating that cranial growth has not yet reached adult levels at P20.

### Surgery

Pups with a visible milk band were removed from the litter and put under isoflurane anesthesia (3-5%; Phoenix Pharmaceuticals, Burlingame, CA). For P16 and P20 pups, the scalp was shaved, and ophthalmic ointment was applied to the eyes. Pups were injected with the anti-inflammatory drug carprofen (5 mg/kg subcutaneously; Putney, Portland, ME) and an incision was made down the midline of the scalp. Using dura scissors, the entire left side of the skull above the neocortex was carefully removed. The brain was flushed as needed with sterile saline to prevent desiccation. A 32-gauge needle was coated with fluorescent DiI (Vybrant DiI Cell-Labeling Solution; Life Technologies, Grand Island, NY) and attached to a stereotaxic arm (Figure 2A). Bregma was defined as the origin of a grid of points on the neocortex, and the needle was inserted into the cortex to a depth of at least 2 mm in a 1 mm grid, including as many locations in the neocortex as the craniotomy allowed (Figure 2B). The needle was reinserted into DiI after three insertions to ensure it remained coated in fluorescent dye.

**Figure 2.**
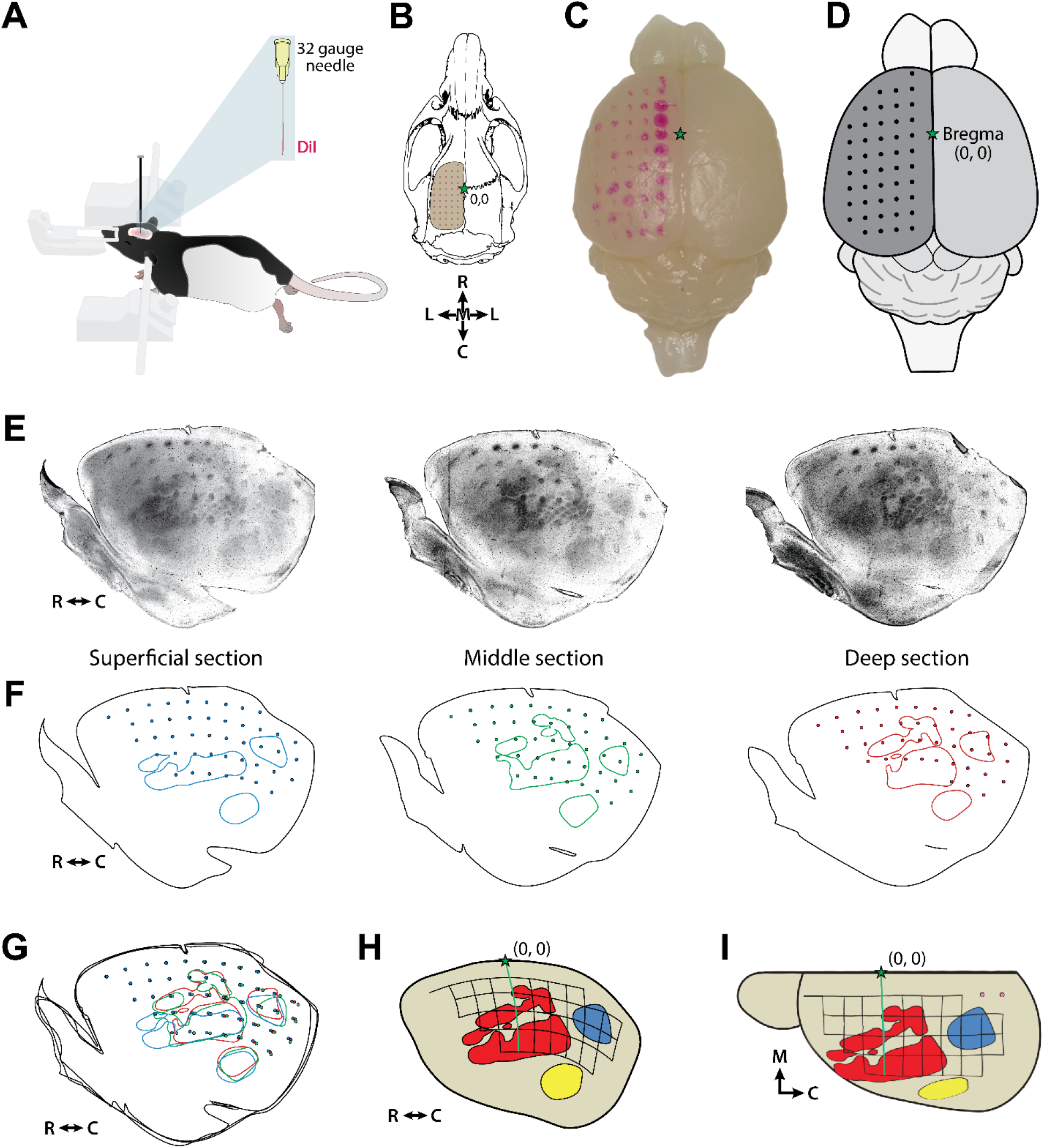
Methodology for stereotaxic coordinate mapping and top-down neocortical reconstruction. **A** Schematic of a rat pup secured in a stereotaxic frame. Inset shows a 32-gauge needle coated with the fluorescent DiI, which is inserted into the neocortex in a grid pattern. **B** Dorsal schematic of the skull with a craniotomy exposing the left cortical hemisphere, showing the stereotaxic grid pattern relative to bregma (0,0; green star). R = rostral, C = caudal, M = medial, L = lateral. **C** Dorsal view photograph of an extracted, post-fixed whole brain displaying the physical grid layout of the fluorescent DiI markers on the surface of the neocortex. **D** Illustration of the brain in **C**, showing the surface coordinate points relative to bregma (0,0). **E** Serial tangential sections of the flattened cortical hemisphere stained for CO. Rostral is to the left, medial is to the top. Panels represent superficial, middle, and deep sections of a representative P12 brain, revealing distinct layer IV boundaries and DiI tissue marks. **F** Corresponding line tracings of the serial sections in **E**, isolating the individual probe locations (dots) and anatomical boundaries (lines) unique to the superficial (blue), middle (green), and deep (red) tissue planes. **G** Composite overlay of three serial sections in **F**, combining all sections into a single comprehensive reconstruction. **H** Top-down reconstruction of the flattened hemisphere, aligning the corrected coordinate grid with defined primary sensory fields (S1, red; V1, blue; A1, yellow) relative to bregma. **I** Final straightened stereotaxic grid template (when viewed from directly overhead) showing coordinate references mapped directly onto the boundaries of the primary cortical areas. Pink-filled dots indicate grid coordinates not seen in the reconstruction.

Following complete mapping of the exposed neocortex, the pup was euthanized with an overdose of 10:1 ketamine/xylazine (>100 mg/kg) and perfused with phosphate-buffered saline (PBS), followed by 4% paraformaldehyde. The brain was immediately extracted (Figure 2C, D) and post-fixed in 4% paraformaldehyde for at least 24 h. At least 24 h before brain sectioning, it was transferred to a 20% solution of sucrose in PBS (PB-sucrose) until it was no longer buoyant in solution.

### Histology

The cortical hemispheres were dissected apart from the underlying tissue, including the hippocampus and basal ganglia, and flattened between glass slides for 5-30 minutes. To prevent over-flattening of the neocortical tissue, 1.5-mm-thick metal spacers were placed on either side of the cortical tissue. Small weights (∼10 g) were used to apply light pressure to the top glass slide. This methodology has been reported previously (Dooley and Blumberg, 2018).

Next, the flattened cortex was placed on a platform of frozen PB-sucrose with the cortical surface facing up, and a glass microscope slide was used to apply light pressure to the cortical hemisphere until it was completely frozen. The flattened cortex was then sectioned tangential to the pial surface at 80 µm using a freezing microtome (Leica Microsystems, Buffalo Grove, IL). Following sectioning, wet-mounted tissue was immediately photographed using light and fluorescent microscopy to identify the relative positioning of fluorescent probes to blood vessels, tissue damage, and any other visual markers.

Sections were then stained for CO, which has been shown in developing rats as young as P5 to reliably delineate primary sensory areas (Seelke et al., 2012). Briefly, cytochrome C (3 mg per 10 mL solution; Sigma-Aldrich), catalase, (2 mg per 10 mL solution; Sigma-Aldrich) and 3,3’-diaminobenzidine tetrahydrochloride (DAB; 5 mg per 10 mL solution; Spectrum, Henderson, NV) were dissolved in a 1:1 dilution of PB-H_2_O and distilled water. Sections were developed in well plates on a rocker at 35–40°C for 3–6 h, after which they were rinsed and mounted. Stained sections were again photographed at 2.5X or 5X magnification, and these individual images were combined into a single composite image.

### Reconstructions

CO staining darkly labels layers IV and VI, with the boundaries of primary cortical areas, particularly the barrels of primary somatosensory cortex (S1) being most striking in layer IV. Thus, our goal with flattening the tissue was to contain layer IV in a single tangential section; however this was often not possible across the entire neocortical hemisphere. Instead, individual sections often contained only partial cortical areal boundaries (Figure 2E, F). However, by photographing the entire series of sections and digitally overlaying them on top of one another (Figure 2G), sequential sections can be combined into a single comprehensive reconstruction (Figure 2H). For each section, the blood vessels, tissue artifacts, fluorescent probes, and architectonic boundaries were used to produce one composite reconstruction. At the lateral portions of the cortex, fluorescent probes begin to resemble lines, rather than distinct points, due to the extreme curvature as the neocortex approaches 90°. When this occurred, the dot reflecting this point on the grid was placed in the most medial location, and the extent of the line of the probe was assumed to be at the same rostral-caudal and medial-lateral position. Using surgical notes, each fluorescent probe was assigned x and y coordinates (relative to bregma).

### Grid-straightening

Because flattening the neocortex deforms the tissue—similar to the deformations inherent in transforming the spherical earth onto a flat map—the composite reconstructions of CO-labeled boundaries were digitally corrected using the free-transform tool in Adobe Photoshop. Images were systematically warped until the reconstructed grid of fluorescent probe marks was restored to an equidistant stereotaxic matrix (Figure 2I).

### Angle tangential to the cortical surface

Because we flattened the neocortex, we were unable to use our tissue to determine the angle of the cortical surface. But this is often an important detail to consider when performing stereotaxic injections, particularly if the objective is to record tangential to the cortical surface. Therefore, we used an existing atlas of developing rat brains, sectioned in the coronal plane, to reconstruct the angle of the neocortex at each point of the grid (Khazipov et al., 2015). These numbers reflected the average of the angle tangential to the cortical surface of the left and right hemispheres for the nearest aged atlas (P8 = P8, P12 = P10, P16 = P14, P20 = P21).

## Results

We reconstructed stereotaxic coordinates over primary cortical areas (S1, A1, and V1) in flattened neocortical tissue for both Sprague-Dawley and Long-Evans rat pups at P8, P12, P16, and P20. Across all ages, DiI-labeled stereotaxic probe tracks were visible in tangential sections spanning the entire cortical depth. CO staining reliably labeled layer IV of primary sensory areas, typically spanning two to three adjacent slices (see Figure 2E), enabling us to draw precise areal boundaries. Areal boundaries and fluorescent probes were readily aligned, creating comprehensive reconstructions of the developing rat neocortex.

At each age, we observed clear, age-dependent shifts in the relative location of S1, A1, and V1. While the general topography remained consistent—S1 lateral and rostral to V1, A1 lateral and caudal to S1—the absolute position of each area relative to bregma shifted with age. Notably, neocortical expansion from P8 to P20 was not uniform: rostral-caudal distances increased much more than medial-lateral distances, although growth was seen in both dimensions. These distortions underscore the limitations of simply scaling adult stereotaxic coordinates down to pup brains.

After aligning CO stained maps with stereotaxic probe locations, we generated warped, age-specific top-down reconstructions for Long-Evans rats (Figure 3) and Sprague-Dawley rats (Figure 4). These maps provide stereotaxic coordinates for primary cortical areas from P8 to P20. Independent reconstructions from the two animals at each age exhibited closely matched structural profiles, confirming that the spatial relationships between primary cortical areas and grid coordinates were conserved. Minor strain differences were observed: Long-Evans rats showed slightly larger and more caudally shifted sensory areas compared to Sprague-Dawley rats at matched ages, likely reflecting differences in skull and brain morphology.

**Figure 3.**
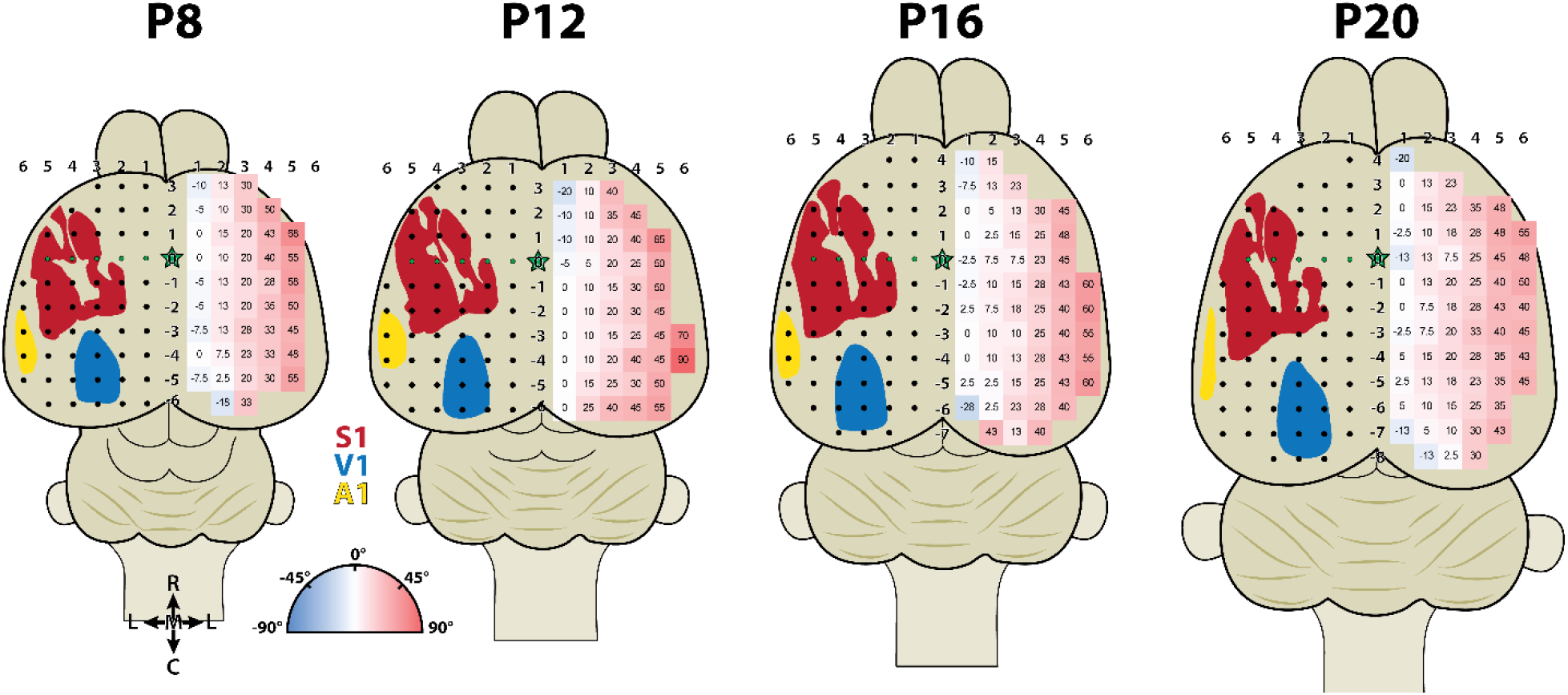
Developmental stereotaxic coordinates and neocortical surface angles in Long-Evans rats. Top-down reconstructions of the neocortex at P8, P12, P16, and P20. On the left neocortical hemisphere, primary sensory areas derived from flattened CO stained tissue boundaries are shown, including the primary somatosensory cortex (S1, red), primary visual cortex (V1, blue), and primary auditory cortex (A1, yellow). Black dots indicate stereotaxic coordinate target locations across the hemisphere, with the green-filled dots indicating bregma (0 mm AP). The green star indicates the location of bregma on the midline. On the right, the local angle of the neocortical surface relative to the horizontal plane (0°) measured at each grid intersection point, taken from Khazipov et al., 2015 (see methods for more details). Numbers inside the grid are the angle tangential to the cortical surface in degrees. Red shading corresponds to positive angles (up to 90°), blue shading corresponds to negative angles (down to -90°), and white indicates a horizontal plane. Medial-lateral grid columns (top numbers) and rostral-caudal grid rows (midline numbers) indicate absolute coordinates in millimeters relative to bregma (0,0).

**Figure 4.**
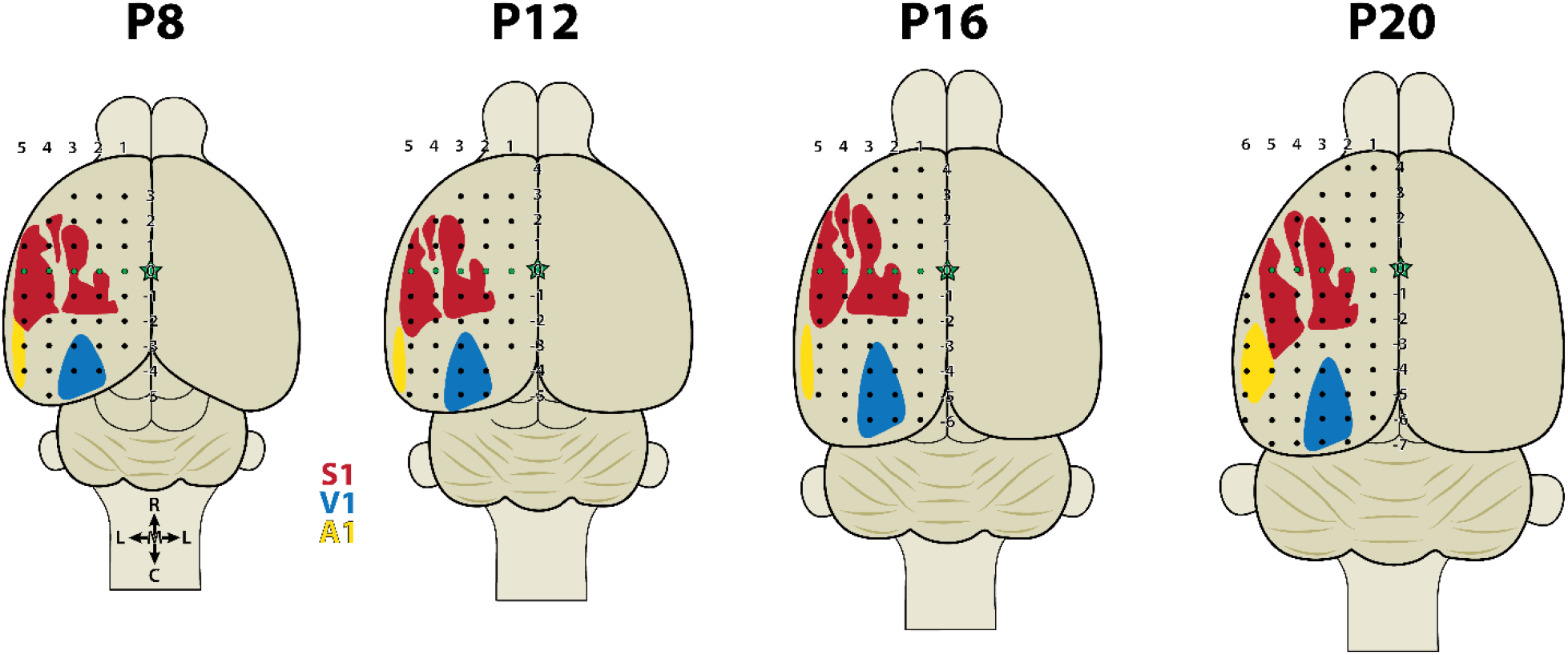
Developmental stereotaxic coordinates in Sprague-Dawley rats. Top-down reconstructions of primary sensory areas (S1, V1, and A1) mapped relative to the stereotaxic grid at P8, P12, P16, and P20. All layout and labeling conventions as in Figure 3, with the exception of the neocortical surface angle maps.

To demonstrate how the neocortical grids aligned onto localized functional subdivisions, we generated high-magnification reconstructions of the S1 barrel field in Long-Evans rats across all four ages (Figure 5). Tangential CO staining clearly resolved individual barrels (rows A1–E1) at P8, with the structural definition of distinct barrel walls distinctly visible at all ages (Figure 5, left panels). Anatomical schematics illustrate the representative spatial relationships between these functional modules and nearby stereotaxic coordinate intersections (Figure 5, right panels). Reflecting the non-linear expansion observed across the hemisphere, the position of the barrel field relative to fixed stereotaxic coordinates shifts systematically across development. For example, the (-1, 4) coordinate target sits near the rostro-medial border of the barrel field at P8 but occupies an internal position near the E1/D1 rows by P16, capturing a local outward and forward displacement of the somatosensory map relative to skull landmarks. Importantly, while the precise boundaries of individual barrels exhibit expected biological variability, the overall topographic progression follows a conserved developmental trajectory, providing a representative reference framework for targeting functional columns within S1.

**Figure 5.**
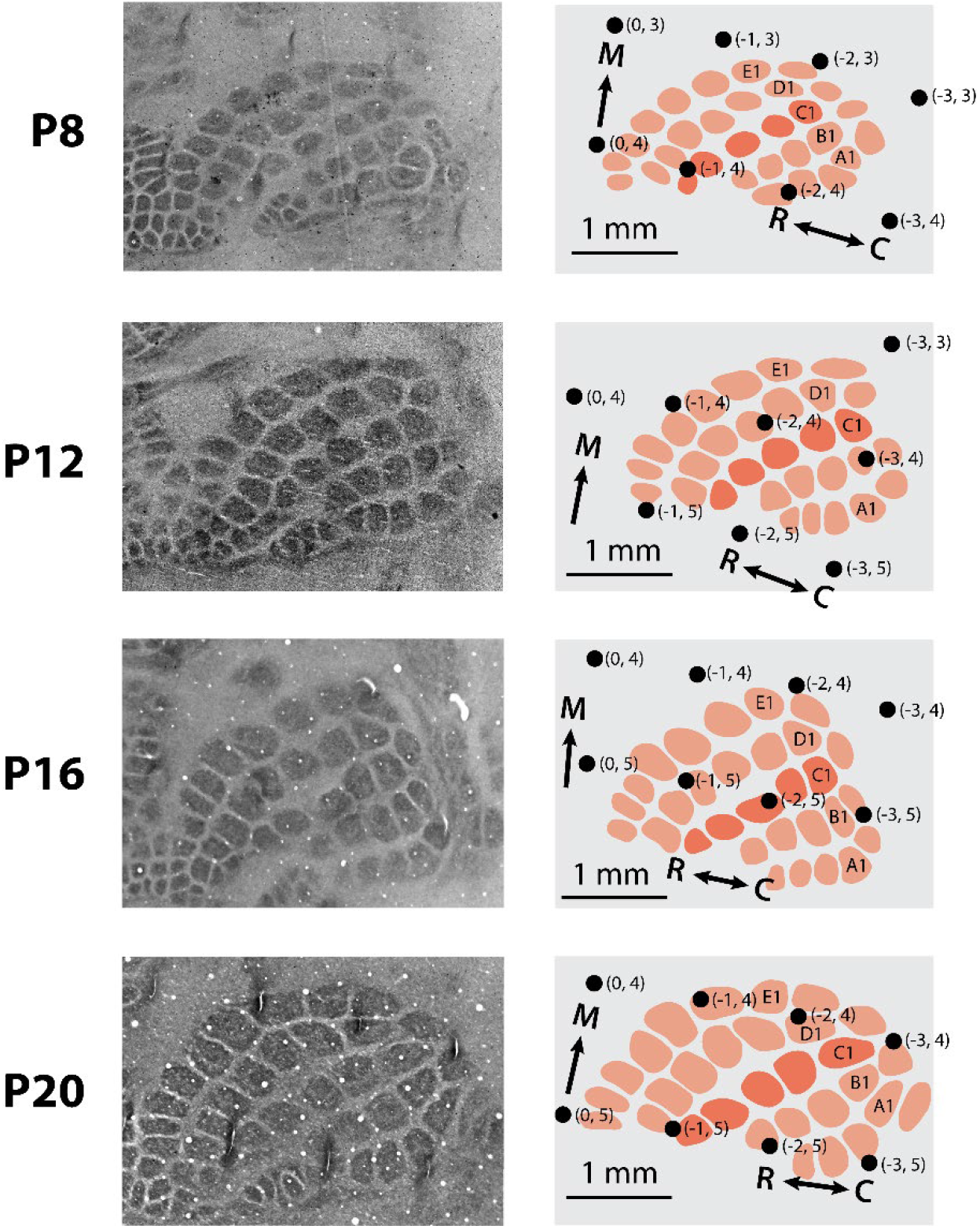
High-magnification developmental tracking of the Long-Evans rat S1 barrel field. Reconstructions track the location of the barrel subfield across four early postnatal ages: P8, P12, P16, and P20. *Left:* Photomicrographs of tangential neocortical sections processed for CO staining, showing the distinct walls of individual barrels. *Right:* Corresponding anatomical illustrations mapping the layout of barrels (red ovals), labeling the first whisker of all rows (A-E). Row C is shown in darker red than other rows. Black dots indicate stereotaxic coordinates with absolute values given in millimeters relative to bregma (R-C, M-L). Scale bars = 1 mm.

## Discussion

The present atlas provides a resource for rat researchers in developmental systems neuroscience by replacing resource-intensive, laboratory-specific trial-and-error coordinate mapping with a standardized, anatomically verified atlas of primary cortical areas. Our results confirm that the growth of the infant rat neocortex from P8 to P20 does not follow a simple, isometric scaling law (Bandeira et al., 2009; Khazipov et al., 2015). Specifically, we find more expansion along the rostral-caudal axis than the medial-lateral axis. By capturing these regional growth distortions and mapping them relative to functional structural boundaries—such as the S1 vibrissae macro-barrels—this atlas provides a reliable blueprint for targeted neurophysiological recordings or viral delivery approaches.

The current age range includes several major functional transitions, including the onset of spatially and temporally precise sensory responses (Colonnese et al., 2010; Dooley and Blumberg, 2018; Dooley et al., 2021; Makarov et al., 2021), behavioral-state-dependent activity (Murata and Colonnese, 2018; Gómez et al., 2021), and eye- and ear-opening (see Colonnese and Phillips, 2018). Importantly, the methods used here are not appropriate at earlier postnatal ages because prior to P5, CO staining fails to resolve distinct areal boundaries (Seelke et al., 2012). However, several developmental atlases exist that focus on the first postnatal week (Khazipov et al., 2015; Ramachandra and Subramanian, 2016) and even prenatal brains (Altman and Bayer, 1995; Paxinos and Ashwell, 2018), relying on different histological techniques to visualize structure within the tissue. And on the other end of development, neocortical and skull growth is not entirely complete at P20; bregma and lambda landmarks have not yet reached adult-like distances in either strain by this stage (see Figure 1B). However, beyond P20, the extreme non-linear growth dynamics characteristic of early postnatal development begin to stabilize. Consequently, standard coordinate scaling based on adult bregma-lambda distances is more likely to yield accurate, predictable stereotaxic results in juvenile rats.

## Acknowledgment

We thank Alexandros Nanos for surgical assistance and John Kobrossi for technical assistance. This research was supported in part by the Sleep Research Society Foundation’s Career Development Award to J.C.D.

